# Identifying an optimal anti-gravity assistance level for select functional shoulder movements: A simulation study

**DOI:** 10.1101/2024.09.25.615099

**Authors:** Morteza Asgari, Dustin L. Crouch

## Abstract

The level of assistance torque is one key design parameter for passive shoulder exoskeletons. High assistance levels may perturb arm movements, while low assistance may not provide functional benefits. This study aimed to use computational tools to identify an optimal anti-gravity assistance level for passive shoulder exoskeletons.

We used the task space framework to perform biomechanical simulations of arm movements in OpenSim (Stanford, CA, USA). The simulated movements included shoulder elevation and lowering movements in frontal and scapular planes, as well as forward and lateral reaching movements. These movements were simulated across a range of assistance torque levels from 0% (no-assistance) to 100% of the maximum shoulder gravity torque, in increments of 10%. The optimal assistance level was identified based on analysis of hand kinematics, muscular response efficiency, and glenohumeral joint stability.

As the assistance level increased from 10% to 40%, the variability of hand movements nearly doubled, and this trend continued for higher assistance levels. The total muscle effort rate was minimized at an assistance level ranging from 20% to 30%. While the stability of the glenohumeral joint was mostly maintained across assistance levels, it decreased slightly at higher assistance levels.

The results of this study indicated that, for the simulated movements, an optimal assistance level lies within the range of 20-30% of the maximum gravity torque at the shoulder joint. Assistance levels above 40% could cause undesired effects such as greater variability of end-limb kinematics, reduced muscular efficiency, and compromised glenohumeral joint stability.

## Introduction

Arm weight compensation has been the design paradigm for many powered and passive (i.e., unpowered) shoulder exoskeletons developed for medical rehabilitation [1-3] and movement assistance/support [4-6]. The shoulder gravity torque generated by the mass of the distal limb segments is much greater than inertial torques during daily activities [7]. This is especially the case for patients with neurological and orthopedic shoulder disability who move their arm at lower speeds than healthy people. Weight compensation of the upper extremity can enhance motor function and extend the reaching workspace for such patients [8]. Thus, many end-effector robots [8, 9], cable-driven manipulators [10] and exoskeletons [11, 12] for clinical applications function as gravity compensators. Gravity compensating exoskeletons have been also developed for motion augmentation and assistance during occupational tasks (e.g., overhead activities or manual handling) [4-6]. Occupational exoskeletons physically aid workers during high-demanding tasks and reduce the risk of work-related injuries by removing excessive musculoskeletal strains and fatigue [6, 13].

Powered upper limb exoskeletons can provide transparent, feedback-controlled, gravity balance in any arm posture [8, 14]. Thanks to the advanced design mechanisms, embedded position/force sensors, and control strategies, powered exoskeletons can provide decoupled, full-arm balance in any pose and trajectory without restricting other degrees of freedom of the arm. However, achieving such flexible gravity-compensating capacity comes with trade-offs, such as high weight, obstruse design, and high maintenance cost, which limit the wearability and portability of powered exoskeletons. These disadvantages have recently given rise to the development of passive exoskeletons as an attractive solution for providing continuous, portable gravity compensation to assist or support arm movements [1, 4, 5, 15].

Passive exoskeletons mainly consist of energy-shuffling mechanisms in which counterweights and/or elastic elements (e.g., mechanical springs and rubber bands) store mechanical energy during specific movement phases and return it during others [4, 16]. Counterweights are avoided for wearable devices since they increase the overall weight of the device. Manufacturing mechanical springs with zero free length or specific force-displacement profiles is also costly [17]. Therefore, multi-linkage [4], parallelogram-[17, 18], pulley-[5], and cam-based [1] mechanisms have been used to modulate the force output of off-the-shelf, non-zero free length springs and, thus, tune the assistance level as desired. Depending on the targeted end user, these mechanisms can help to produce arm balance [17, 18] at certain angles or modulated bionic assistive torque across the range of arm elevation [1, 4, 6].

The assistance level is a crucial design parameter for passive shoulder exoskeletons. However, there is no consensus on what assistance level is most beneficial to the user [5]. A study of the ShoulderX exoskeleton [19] reported up to 80% reduction in activation of the anterior deltoid and upper trapezius muscles with an assistive torque ranging from 8.5 to 20.0 Nm. In an experimental evaluation of the Proto-MATE [6] with an assistive torque equal to 50% of the estimated gravity torque (i.e., up to 5.5 Nm), activations of the shoulder flexor muscles were reduced by up to 43%. Rossini *et. al*. [5] also reported up to 22% reduced activation of shoulder flexor muscles when the Exo4Work shoulder exoskeleton delivered 3 Nm assistive torque. However, in a study with able-bodied subjects, Asgari et. al. reported that users experienced discomfort [15] and greater muscle activity during arm lowering movements when a wearable passive shoulder exoskeleton compensated for 50% of the shoulder gravity torque [1]. Some studies also suggest that an optimal level of assistance might be task dependent [28].

Increasing assistance level can substantially reduce the activity of the muscles responsible for positive shoulder elevation/flexion against gravity. However, too much assistance can be associated with undesirable effects such as higher levels of activation in antagonist muscles [19-21], compromised joint stability and stiffness [22, 23], and altered joint kinematics and end-point precision [24]. Studies of occupational exoskeletons reported that, during overhead reaching tasks, the activity of shoulder flexor muscles (e.g., deltoids, trapezius) was less, while the activity of antagonist muscles (e.g., triceps and latissimus dorsi) was greater, with the exoskeletons than without [19, 21]. When faced with external assistive forces or perturbations, the neuromuscular system stabilizes the upper body joints and end-limb by modulating the magnitude and patterns of activations of agonist-antagonist muscle pairs [25, 26]; this has important implications for joint stiffness and stability. Specifically, over-assistance could compromise agonist-antagonist muscle coordination and joint and end-limb stability [27, 28], resulting in higher muscle activations and reduced task performance and precision for the wearer [20].

The main objective of this simulation-based sensitivity study was to determine the effect of anti-gravity assistance level at the shoulder on (1) muscular response efficiency and (2) stability at the joint and end-limb. Our hypothesis was that there exists an optimal assistance level, above which may degrade GH joint stability or task performance, and below which may provide less improvement in muscular efficiency. Our overall approach was, using a task space (i.e., operational space) framework, to perform predictive biomechanical simulations of selected arm reaching and elevation movements; simulations were performed for eleven different anti-gravity assistance levels ranging from no assistance (0%) to 100% of the maximum gravity torque at the shoulder, in increments of 10%.

## Methods

### The musculoskeletal model

For our musculoskeletal simulations, we used an existing upper limb musculoskeletal model implemented in OpenSim (version 4.1) [29]. The upper extremity model includes 7 degrees of freedom (DOFs), with sets of three, two, and two DOFs describing the shoulder, elbow, and wrist kinematics, respectively. The skeletal geometry of the model is defined based on the anthropometric data of a 50^th^ percentile male. The model is actuated by fifty Hill-type muscle-tendon units crossing shoulder, elbow, and wrist joints.

### The task space framework

The task space (i.e., operational space) framework is a model-based control approach to track a desired trajectory defined in the task coordinate system [30-34]. Unlike the joint coordinate systems, the task coordinate system is directly attached to the end-effector whose motion and contact forces describe the performed task. Therefore, developing the equations of motion in the task coordinate system simplifies the control problem by abstracting away the generalized coordinates in the joint space. The task space controller incorporates a dynamic model of the control plant to directly estimate the joint torques that are required to execute a dexterous and compliant tracking motion (i.e., task). This control strategy resembles the way humans control their upper limb motion where the central nervous system (CNS) acts as a high-level controller and executes goal-oriented hand movements. Among different task space controllers designed for over-actuated systems (i.e., redundant), Khatib’s [30, 35] approach appealing as it decouples the dynamics of the task from that of the null space (i.e., posture). This approach is well-suited for musculoskeletal models, such as the upper limb model used in this study, in which the number of DOFs, *n*, is larger than the task dimension, *m* [35, 36]. Since musculoskeletal models are also subject to different constraints *Φ*(*q*) = 0 (e.g., holonomic), we used an orthogonal projection scheme, proposed by Aghili [37, 38], to firstly eliminate the constraint forces by projecting the plant dynamics onto the constraints’ manifold.

The joint-space equation of motion (EOM) for the upper limb with *n* DOF and *k* linearly independent constraints, whose generalized coordinate vector is *q ∈ ℝ*^*n*^, can be written as [33]:

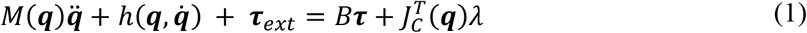

where *M*(q) *∈ ℝ*^*n*×*n*^ is the mass matrix; 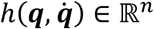 ) *∈ ℝ*^*n*^ includes gravity, centripetal, and Coriolis forces; τ_*ext*_ *∈ ℝ*^*n*^ is the vector of externally imposed forces/torques; τ *∈ ℝ*^*n*^ is the vector of generalized forces; *B* = *diag*(*b*_1_, … , *b*_*n*_) with *b*_*i*_ = 1 for actuated coordinates and *b*_*i*_ = 0 for passive coordinates; 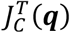 is the transpose of the constraint Jacobian; and λ *∈ ℝ*^*k*^shows the constraint forces. For the sake of brevity, we replace *M*(q) by *M, h*(q, 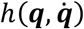 ) by *h* from this point forward.

To eliminate the constraint forces form (1), we multiply both sides by an orthogonal projection operator *P* (*P* is orthogonal if *P* = *P*^2^ = *P*^*T*^) to obtain the projected EOM [37, 39]:

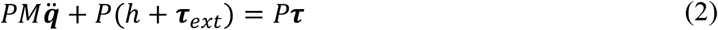

Since 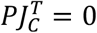 , we can obtain 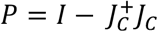 , where 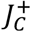 is the Moore-Penrose pseudoinverse of the constraint Jacobian *J*_*C*_ . We can conclude from the first differentiation of the *k*constraint 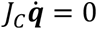 that the vector of generalized velocity lies in the null space of the constraint Jacobian matrix [37]. This concludes:

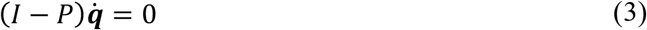

Differentiating form (3):

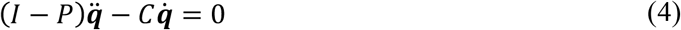

and 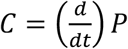 and can be obtained as 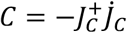. Substituting (4) in (2) gives:

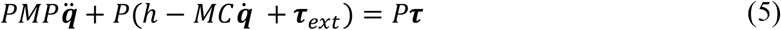

Multiplying (4) by (*I − P*)*M* and adding it to (5) yields:

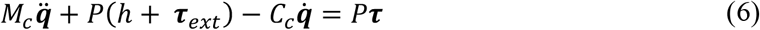

where *M*_*c*_ = *PMP* + (*I − P*)*M*(*I − P*), and *C*_*c*_ = (*I −* 2*P*)*MC*. Unlike the *PM* term in (2), *M*_*c*_ is always invertible and symmetric, preserving the physical characteristics of the inertia matrix *M* [37]. To map the projected joint-space EOM onto the task space, we multiply (6) by 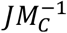 and substitute 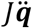 with 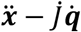 [30, 33]. Note that every task *x ∈ ℝ*^*m*^ with *m* DOFs can be related to the generalized coordinates by the task Jacobian 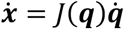.

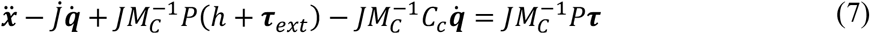

The joint-space generalized forces, τ, can also be similarly mapped onto the task space forces, *F*, via τ = *J*^*T*^*F*. Thus, the EOM can be defined in the task space:

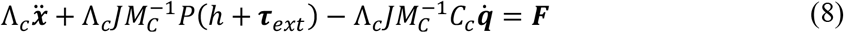

where 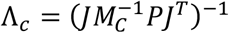 and is called a constraint-compliant inertia matrix. In the case where τ_*ext*_ is available, as outlined by Khatib [30], the controller law can be modeled as:

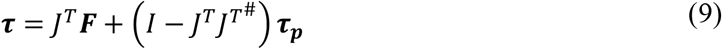

where 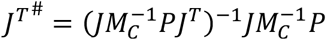 , and τ _*P*_ *∈ ℝ*^*n*^ is the posture vector defining the joint impedance behavior. 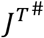 guarantees the decoupling of the task space from the posture space (i.e., the null space). In equation (9), *F* can be computed from (8) based on a command task acceleration 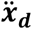 that defines the desired trajectory for the end-limb [30, 33, 34]:

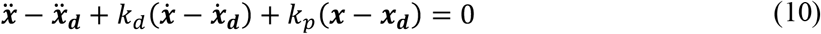

where *K*_*d*_ and *K*_*p*_ are positive-definitive gain matrices and *x* = *f*(q) is the forward kinematics tracking trajectory (Figure 1). In a similar way, τ_*P*_ can be estimated via a command joint acceleration 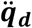 that defines the impedance behavior in the null space [40]:

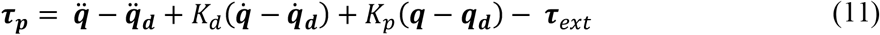

**Figure 1.**
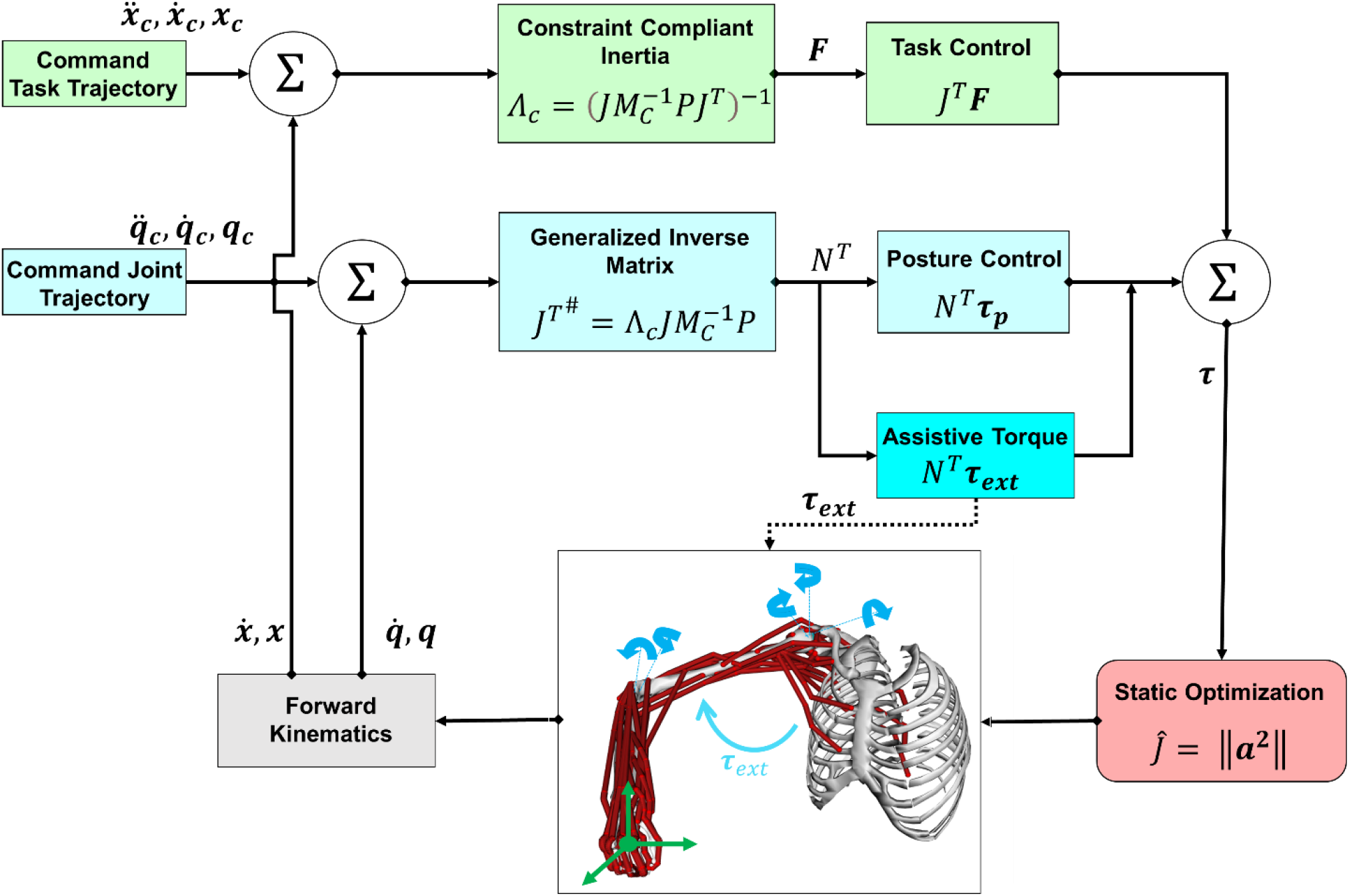
A task/posture space upper limb controller with static optimization to estimate muscle activations.

*K*_*d*_ and *K*_*p*_ are positive-definitive matrices representing, in turn, the damping and stiffness.

### Estimation of muscle activations

We used static optimization (SO) to address the muscle redundancy problem [36, 41]. SO estimates the optimal muscle activations *a*_*i*_ such that muscle forces produce the generalized torques specified by the task space controller, i.e., equation (9), while minimizing the following performance criterion:

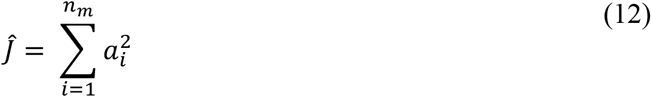

subject to:

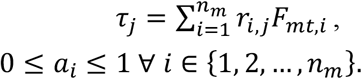

where *r*_*i*,*j*_ is the moment arm of muscle *i* about the axis joint *j*, and *F*_*mt*,*i*_ represents the force generated by the musculotendon unit *i*. We assumed a linear relationship between muscle activation and muscle force and ignored the muscle activation and contraction dynamics.

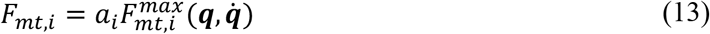

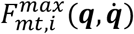 represents the maximum force generating capacity for the *i*^*th*^ muscle estimated based on the instantaneous generalized coordinates and speeds [41].

### Simulations

The presented task space framework was used to execute predictive upper limb movement simulations with eleven levels of assistance at the shoulder joint for select functional tasks. The targeted assistance levels *k*are no-assistance (k=0) and 10, 20, 30, 40, 50, 60, 70, 80, 90, and 100, with *k*being a percentage of the maximum gravity torque at the shoulder. Using the upper limb model, we estimated the maximum gravity torque in the frontal plane with a fully extended elbow and 90-degree arm elevation as following:

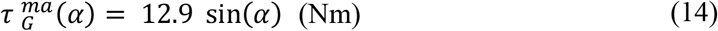

where τ_*G*_ (*α*) is the gravity torque and *α* is the shoulder elevation angle [1, 15, 29]. For simplification, we used an idealized coordinate actuator to model the external assistive torque in the positive direction of the shoulder elevation coordinate. The assistive torque was modeled by the following equation:

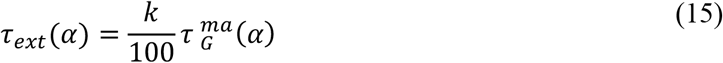

The select functional tasks included shoulder elevation and lowering movements in two different planes of elevation and reaching movements to four target positions (Figure 2). We selected these reaching and elevation movements as they are clinically significant and involve complex coordination of multiple upper joints [42]. Furthermore, these motions require positive/negative elevation, which is a significant movement of the shoulder during daily activities[43].

**Figure 2.**
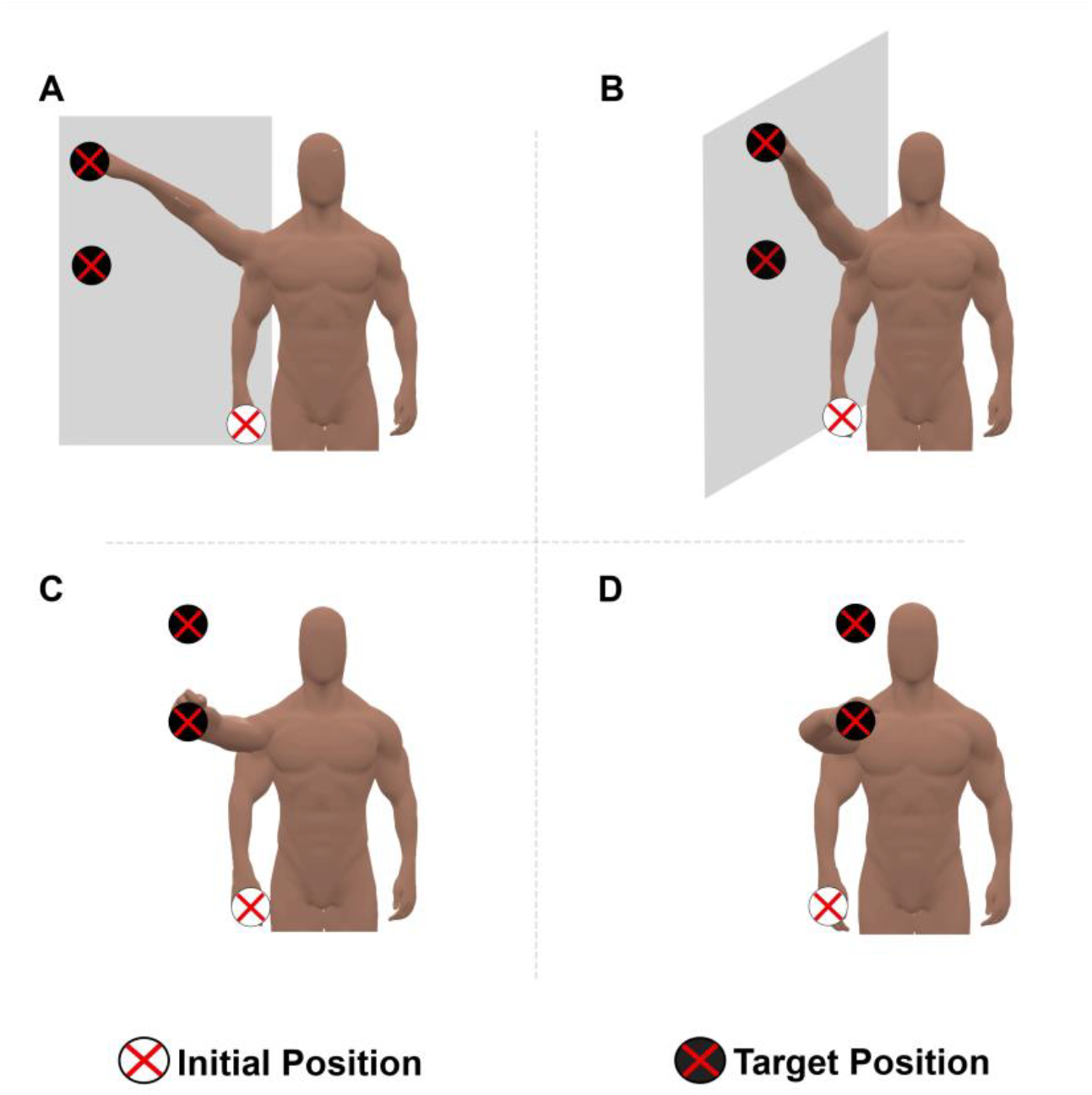
Select functional shoulder movements. Shoulder elevation in the coronal plane (A), and scapular plane (B), lateral reaching (C), and forward reaching (D) movements. Simulations were performed at two targets positions for each of the above movements. For the elevation movements, targets were placed at the shoulder height and above shoulder height such that they can be reached by fully extended elbow. For the lateral and forward reaching movements, targets were closer to the body and can be reached with flexed elbow.

The shoulder elevation movements were simulated in the coronal (i.e., plane of elevation 0º, Figure 2(A)), and scapular planes (i.e., plane of elevation 30º, Figure 2(B)) for the two targets, one placed at shoulder height and one above shoulder height. The target positions during elevation were selected such that they can be reached with the elbow extended. For the lateral (Figure 2 (C)) and forward reaching (Figure 2 (D)) movements, the hand moved from an initial position alongside the body to target positions in front of the model at above shoulder- and shoulder-height levels. For the forward reaching movements, the above-shoulder target was located 25 cm superior and 45 cm anterior to the GH joint center, and shoulder-height target was located 50 cm anterior and at the same height as GH joint center. For the lateral reaching movements, the targets were shifted 15 cm lateral to the GH joint.

The movement planning for the simulated functional tasks was encoded based on equation (10), where the hand’s center of mass (i.e., end-limb) tracks the desired positional trajectory kinematics with three DOFs as the command input. We locked the wrist joint during all the simulations while the other five DOFs were set free, guaranteeing kinematic redundancy. Table 1 includes the initial angles as well as the lock/unlock status of the model’s DOFs during the simulated movements.

**Table 1.**
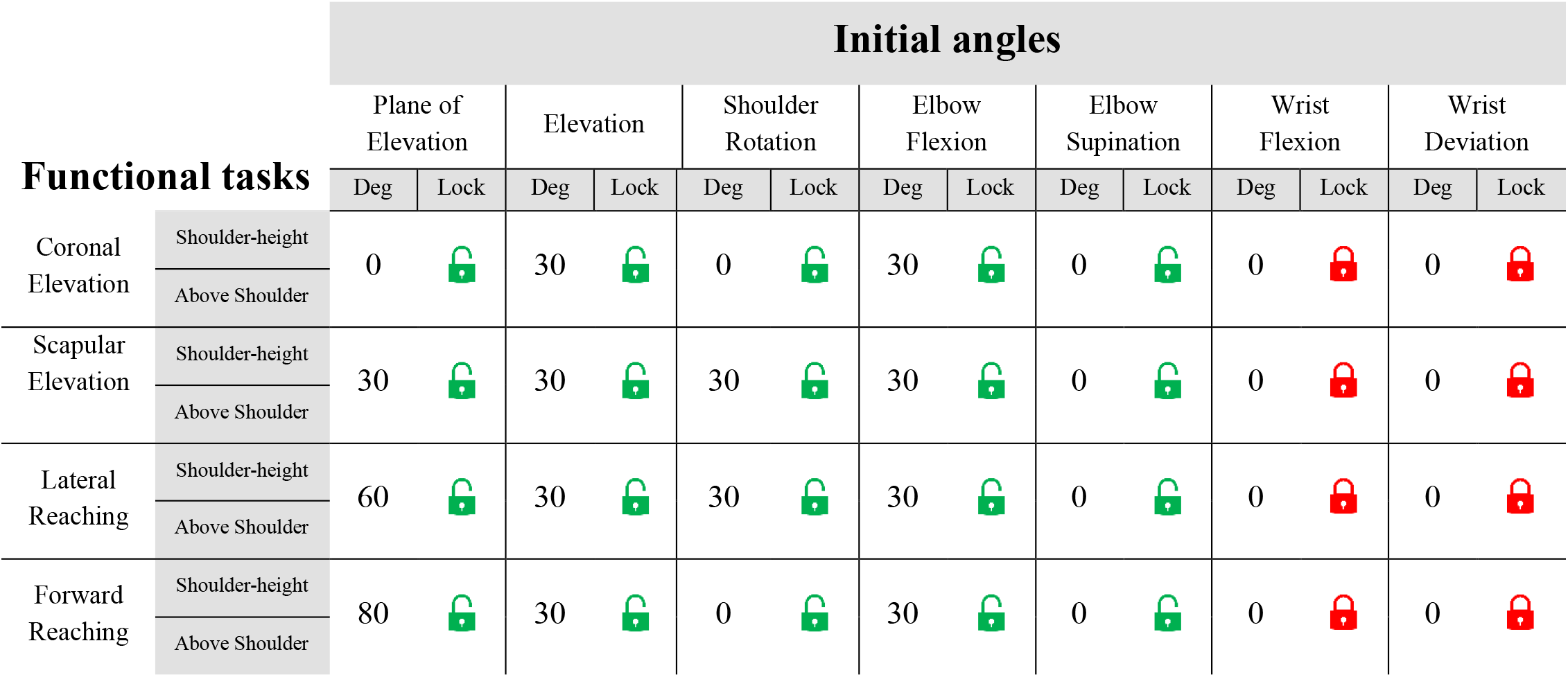
Initial angles and the joint lock/unlock status for the simulated functional tasks.

| Functional tasks |  | Initial angles |  |  |  |  |  |  |  |  |  |  |  |  |  |
| --- | --- | --- | --- | --- | --- | --- | --- | --- | --- | --- | --- | --- | --- | --- | --- |
|  |  | Plane of Elevation |  | Elevation |  | Shoulder Rotation |  | Elbow Flexion |  | Elbow Supination |  | Wrist Flexion |  | Wrist Deviation |  |
|  |  | Deg | Lock | Deg | Lock | Deg | Lock | Deg | Lock | Deg | Lock | Deg | Lock | Deg | Lock |
| Coronal Elevation | Shoulder-height | 0 |  | 30 |  | 0 |  | 30 |  | 0 |  | 0 |  | 0 |  |
|  | Above Shoulder |  |  |  |  |  |  |  |  |  |  |  |  |  |  |
| Scapular Elevation | Shoulder-height | 30 |  | 30 |  | 30 |  | 30 |  | 0 |  | 0 |  | 0 |  |
|  | Above Shoulder |  |  |  |  |  |  |  |  |  |  |  |  |  |  |
| Lateral Reaching | Shoulder-height | 60 |  | 30 |  | 30 |  | 30 |  | 0 |  | 0 |  | 0 |  |
|  | Above Shoulder |  |  |  |  |  |  |  |  |  |  |  |  |  |  |
| Forward Reaching | Shoulder-height | 80 |  | 30 |  | 0 |  | 30 |  | 0 |  | 0 |  | 0 |  |
|  | Above Shoulder |  |  |  |  |  |  |  |  |  |  |  |  |  |  |

The hand’s desired trajectory was decoded based on a fifth-order polynomial function to minimize jerk between the initial and target positions . Such a function also produces bell-shaped velocity profiles that are commonly observed for voluntary point-to-point arm movements. The derivative and proportional gain matrices were set to *K*_*p*_ = 100 and *K*_*d*_ = 20. We did not consider a command joint acceleration for the null space control as in equation (11) and set 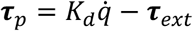 with *K*_*d*_ = *−*0.2*5* . The external torque τ_*ext*_ is the modeled gravity-compensating assistive torque, which was estimated using equation (15). Each of the select functional tasks consists of advancement and retreat phases. During the advancement phase, the hand moves from its initial position to a target position while during the retreat phase it returns to the initial position. The duration of each simulated task was estimated by dividing the hand-to-target distance over the mean target-approaching velocity for the able-bodied participants reported in [42]. The advancement and retract phases had the same execution time, with a 500-millisecond interval between them when the hand was held at the target.

### Data analysis

#### Hand Kinematics

The proposed task-posture controller assumes that the control of the assistive torque takes place in the posture space (i.e., null space) of the task, where the acceleration generated by the assistive torque should not interfere with the acceleration of the hand positional task. Thus, we tracked the position of the hand’s center-of-mass (COM) in 3-dimensional (3-D) space during the simulations. Then, for each assistance level, we calculated the variability in hand position across the eight simulated tasks. To calculate the hand position variability, we resampled the simulations with different assistance levels to the same number of points for every task, and used following equation:

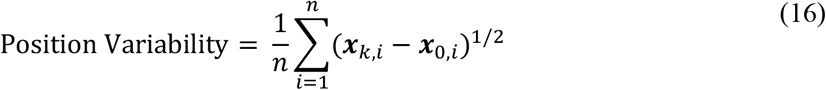

where *k* and *i* represent the assistance level and resampled point number, respectively. *n* is the total number of resampled points for every task, and 0 subscript stands for the no-assistance level simulations. It is expected that the hand position variability should not increase for an optimal level of assistance.

#### Muscular Response Efficiency

During dynamic or quasi-static tasks, the involved muscles can have a load-bearing or fine-tuning contribution depending on whether they lengthen or shorten [27]. To compare the muscular activity across the assistance levels, we computed the total muscle effort rate and efficiency. The total muscle effort rate, ^Ė^_*total*_, was defined as the sum of the individual muscle output calculated based on equation (17).

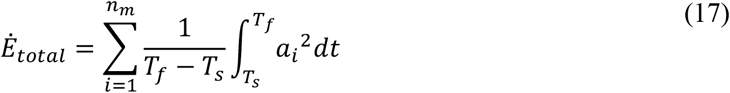

*T*_*s*_ and *T*_*f*_ are the start and finish times of each simulated task, *a*_*i*_ is the muscle activation estimated in (13), and *n*_*m*_ is the number of shoulder muscles. The included shoulder muscles were anterior, middle, posterior deltoids, supraspinatus, infraspinatus, subscapularis, teres major and minor, pectoralis major, latissimus dorsi, coracobrachialis, triceps brachii, and biceps brachii. To quantify the muscular response efficiency, we first categorized the shoulder muscles as being either agonist or antagonist during each phase of the simulated tasks based on whether their fiber length was shortening or lengthening. The categorization was performed for the no-assistance trials and assumed to stay the same across assistance levels. Our anticipation is that the gravity-compensating assistive torque will result in decreased activation of both agonist and antagonist muscles, as it aligns with the purpose of these muscles to generate torque in order to counteract the gravity force. However, the decreasing trend for the antagonist muscles is expected to be limited since they have a fine-tuning function to provide stiffness for the shoulder joint and decelerate the arm. Therefore, depending on the task type, with an optimal level of assistive torque, the synergistic function of agnostic-antagonist muscles is expected to be most efficient, indicated by a maximum reduction in their activations. Therefore, the efficiency was defined as:

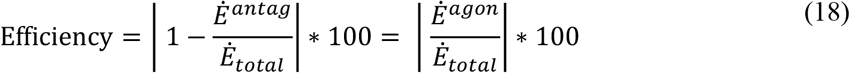

where Ė^*antag*^ and Ė^*agon*^are the total muscle effort rate for the antagonistic and agonistic shoulder muscles, respectively.

#### GH stability

We used the joint reaction analysis tool in OpenSim to estimate the GH joint contact forces [44, 45] during the simulated tasks. To determine the spatial orientation of the glenoid fossa, we placed extra markers around the center of the glenoid fossa in the musculoskeletal model and used their cartesian coordinates to construct a local cartesian coordinate system aligned with and centered at the glenoid rim. This enabled us to transform the estimated contact forces to the anatomical coordinate system of the glenoid fossa (Figure 3) at each time instant and compute the compressive and shear forces at the GH joint. The compressive component pushes the humeral head into the glenoid concavity while the shear components in the superior-inferior (SI) and anterior-posterior (AP) directions destabilize the joint by causing the humeral head to translate away from the center of the glenoid fossa [46]. In this study, we calculated the ratio of SI and AP shear forces to the compressive force as *θ*_*SI*_ and *ϕ*_*AP*_ , respectively, and quantified the GH joint stability as:

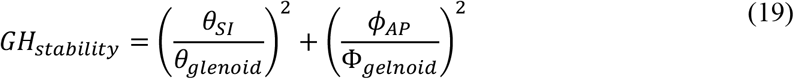

**Figure 3.**
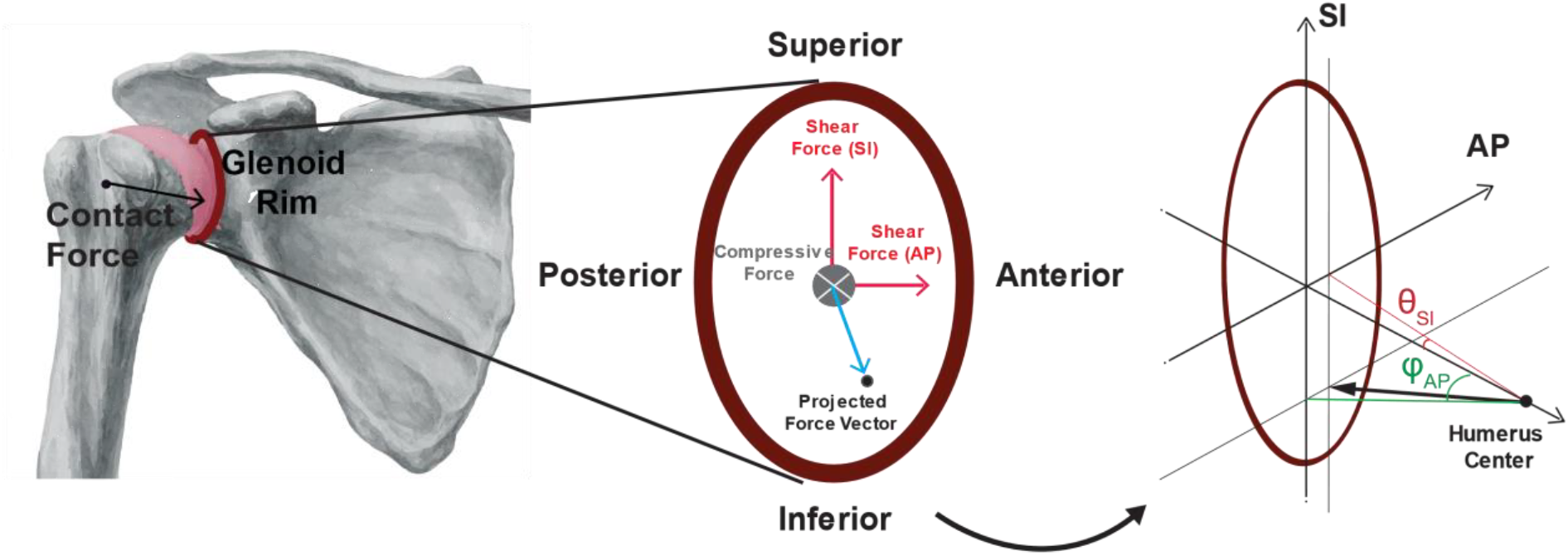
Glenohumeral joint stability. Contact forces at GH joint were transformed to the glenoid fossa, which is represented as an ellipse, and decomposed into compressive and shear forces in superior-inferior (SI) and anterior-posterior (AP) directions. The ratio of SI and AP shear forces to the compressive force were then calculated as *θ*_*SI*_ and *ϕ*_*AP*_ angles and used to determine the GH joint stability.

Equation (19) estimates the glenoid rim as an ellipse whose width and height are 28.4 mm in the AP direction and 36.6 mm in the SI direction, respectively [47]. *θ*_*glenoid*_ and *Ф*_*gelnoid*_ are the maximum angles that the GH joint contact force can have in the SI and AP direction, respectively. For *GH*_*stability*_ *>* 1, when the resultant GH joint contact force points outside of the glenoid rim, the GH joint is considered unstable; conversely, for *GH*_*stability*_ *<* 1 , when the resultant GH joint contact force points within the glenoid rim, the GH joint is considered stable. The *GH*_*stability*_ was calculated at each sampling point for all the tasks.

## Results

### Hand Kinematics

The level of assistance directly affected the variability of the hand position across the simulated tasks. Figure 4 shows an example of the components of the hand trajectory for coronal elevation to a target placed above shoulder-height for the no-assistance and 40% assistance trials. Across all tasks, as the assistance level increased from 10% to 40%, the hand variability almost doubled and surpassed 10 cm (Figure 5). The variability continued to increase beyond an assistance level of 40%, indicating dynamic instability for the hand in the simulated shoulder tasks. Dynamic instability can be defined as the inability of the upper limb model to closely track the desired hand trajectory resulting in lack of precision in task execution [48]. Figure 6 further highlights this instability, showing that the resultant hand COM trajectory deviates from the desired trajectory for assistance levels beyond 40%.

**Figure 4.**
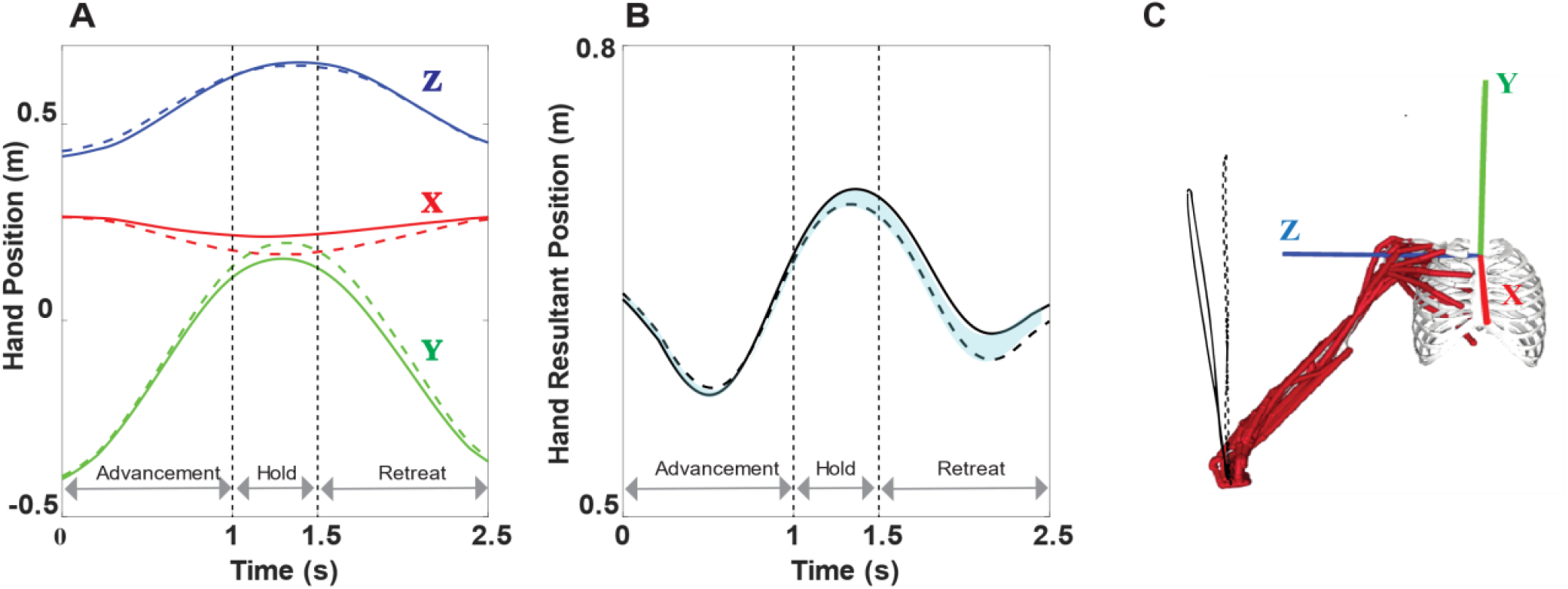
Hand position variability. (A) Hand COM trajectory components for the no-assistance (solid lines) and 40% assistance (dashed lines) trials when shoulder elevation to the above-shoulder target is simulated in the coronal plane. (B) The shaded area between the solid and dashed curves shows the resultant deviations across the movement cycles for the no-assistance and 40% assistance simulations. (C) The musculoskeletal model and 3D trajectory of the hand COM for the no-assistance (solid curve) and 40% assistance (dashed curve) simulations.

**Figure 5.**
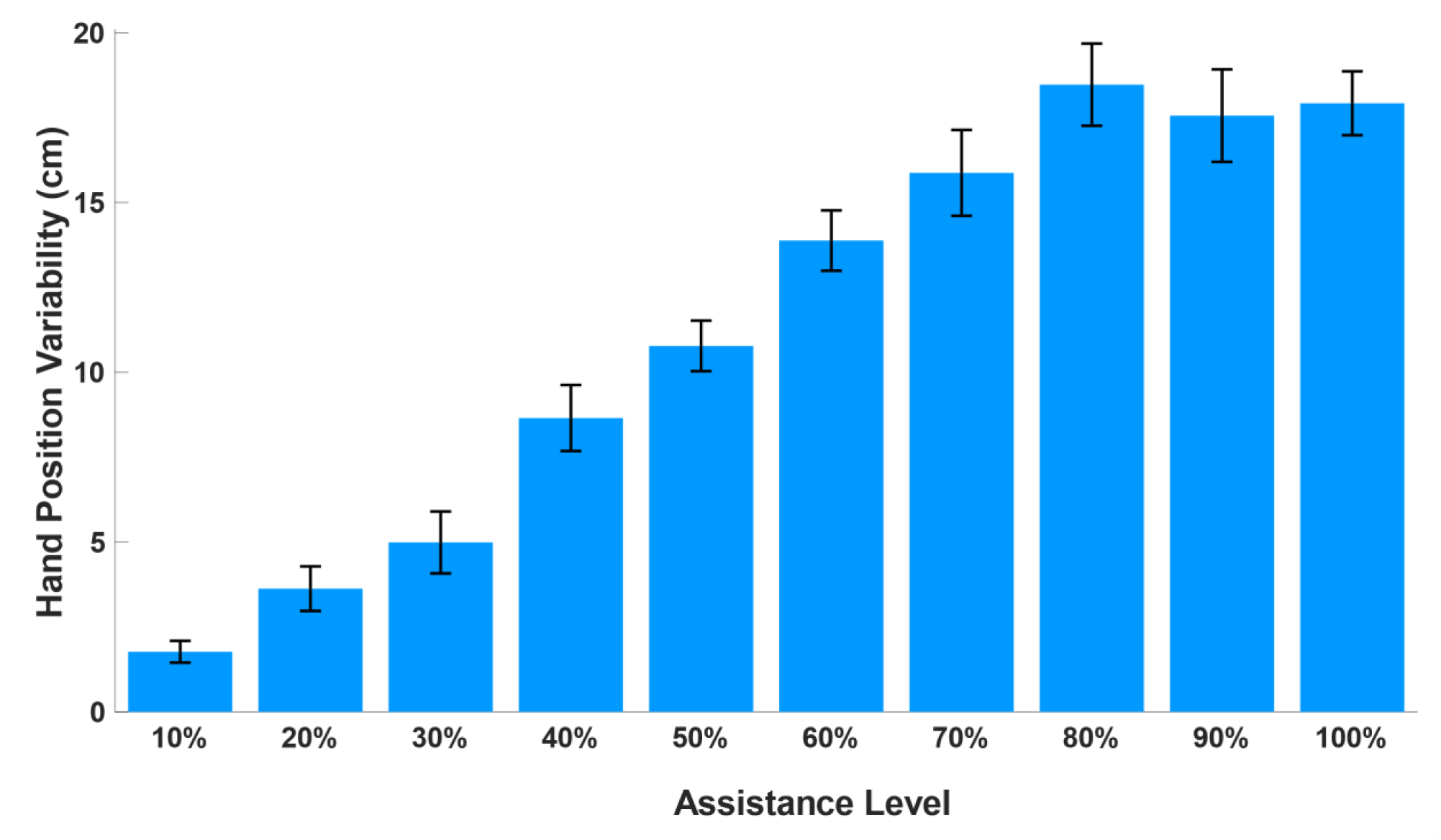
Hand position variability vs. assistance level for the simulated shoulder tasks. The bars and error bars show the mean the standard error of the end-point variability across the shoulder tasks.

**Figure 6.**
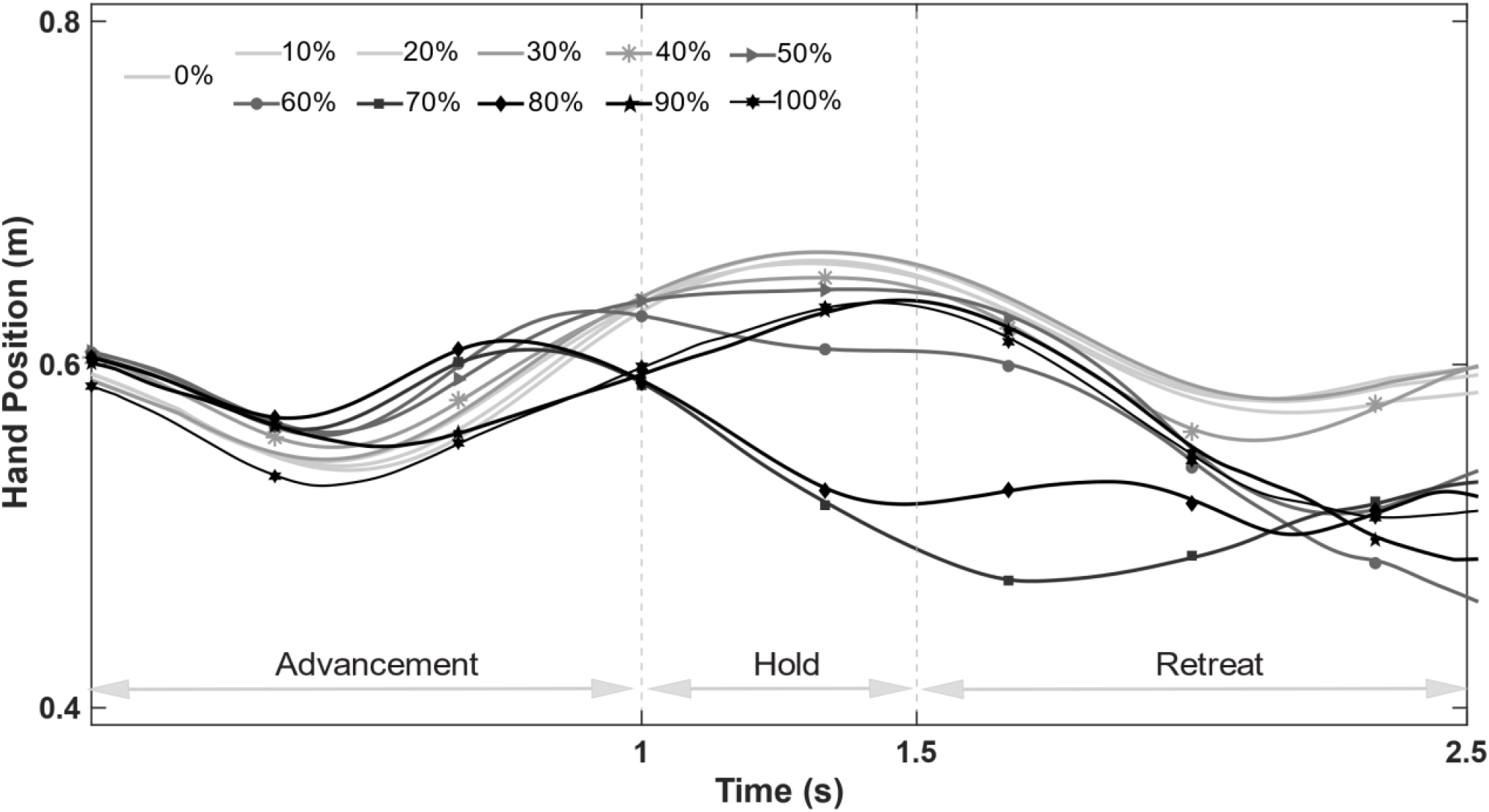
Hand resultant position for the scapular elevation to an above shoulder target. For assistance levels above 40%, the hand COM trajectory shows higher divergence, less precision, and dynamic instability.

The model performed well in terms of following the desired trajectory during the acceleration phase at assistance levels below 50% (Figure 6). However, as the phase progressed to the deceleration part, there was an increasing deviation from the desired trajectory. At assistance levels of 50% and higher, the model exhibited higher divergence from the desired trajectory, resulting in an unstable hand trajectory for the entire simulation. It’s worth noting that the initial position of the hand COM moved further away from the body when the assistance level was set at 50% and higher.

### Muscular Response Efficiency

The level of assistance torque affected the total effort rate during simulated tasks. In general, an initial decrease in the total effort rate was observed as the level of assistance increased, reaching a minimum at an assistance level around 20%-30% (Figure 7). As the assistance level increased beyond 30%, the total effort became larger. Notably, the lateral and forward reaching movements to the shoulder level target deviated from this trend, as their total effort rate increased when the assistance level exceeded 10%.

**Figure 7.**
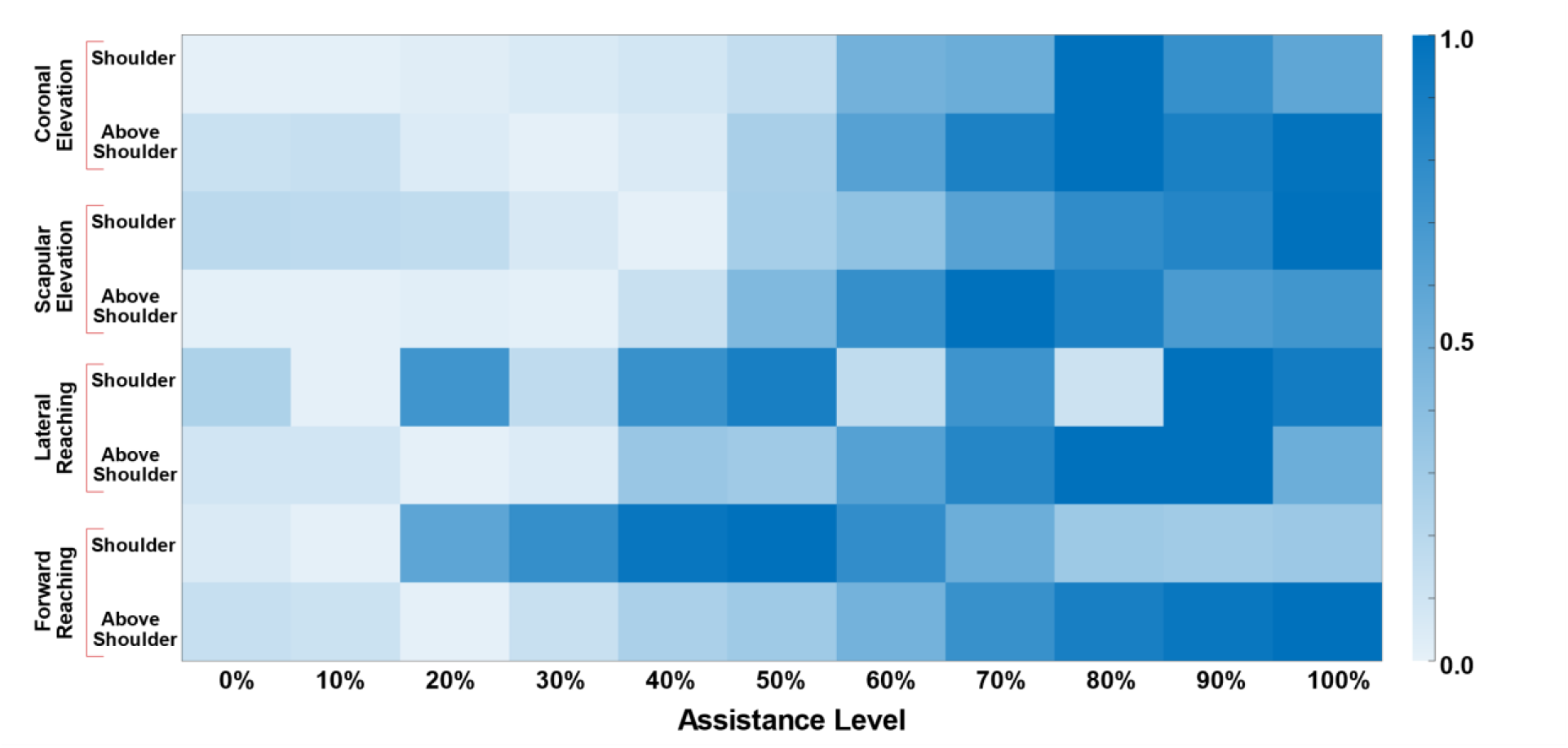
Total muscular effort rate for the simulated shoulder tasks. The sum of the effort rate of the shoulder muscles across the assistance levels was scaled to [0, 1]. Light color cells show less effort rate and darker ones show higher effort rate for the simulated task.

Efficiency, as defined in (18), ranged from 45% to 57% across all assistance levels (Figure 8). A large decrease in efficiency was observed for assistance levels greater than 20% compared to the no-assistance condition. Specifically, the average efficiency decreased from 54.10% to 51.81% with an increase in assistance torque from 20% to 30%. However, for assistance levels higher than 30%, efficiency remained consistently below 54.11%, without a clear trend.

**Figure 8.**
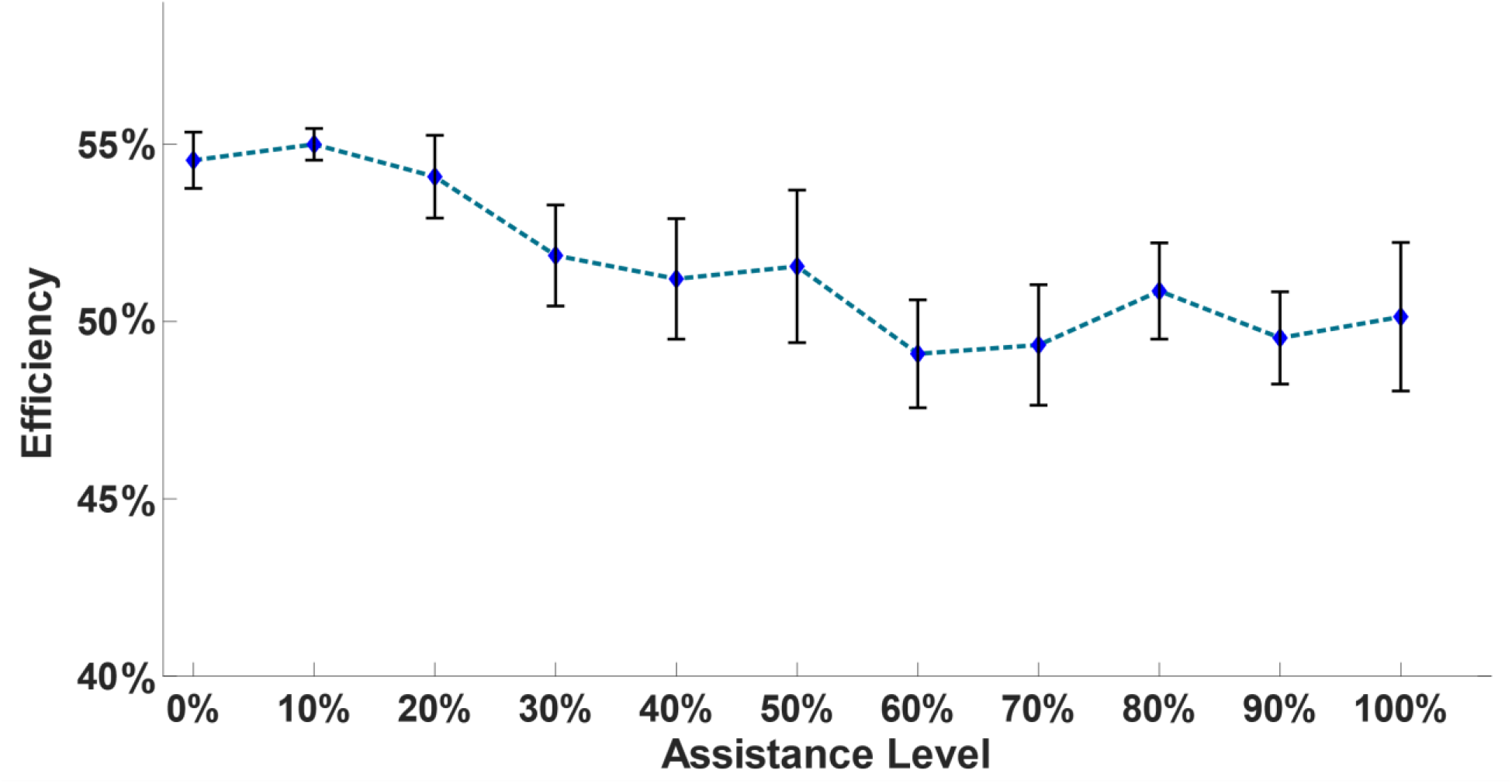
Mean efficiency ± standard error across the simulated task. The estimated efficiency reduces when assistance level is higher than 20%.

### GH stability

As can be seen in Figure 9, the median and variability of *GH*_*stability*_ remained below one across all assistance levels. This suggests that the stability of the GH joint was mostly maintained across simulations, and that the resulting contact force at the GH joint was oriented within the glenoid rim. Between assistance levels of 10% and 30%, the median showed a slight decrease, while the interquartile and upper quartile ranges and the peak of the upper quartile increased compared to the no-assistance condition. At assistance levels of 40% or higher, the median and the magnitude and number of outliers generally increased with assistance level.

**Figure 9.**
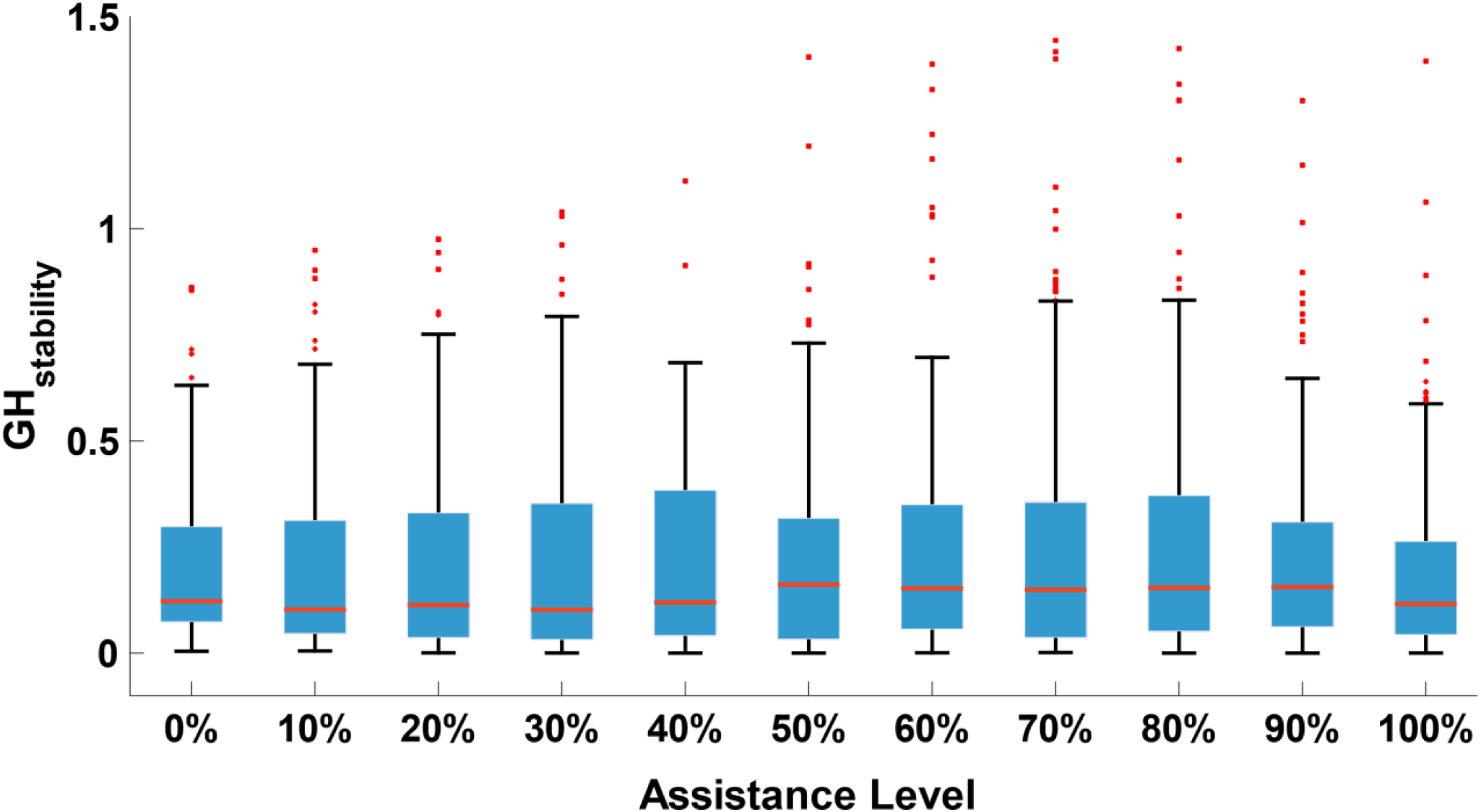
Box plot showing glenohumeral stability index vs. assistance level. The error bars show the standard error within the simulated tasks. For GH_Stability_ < 1, the GH joint is considered stable.

Figure 10 depicts the overlay of the AP and SI components of GH joint contact forces on the glenoid fossa for the lateral reaching task as an example. Consistent with the findings presented in Figure 9, the projected points onto the glenoid rim spread over a larger trace as the assistance level increases up to 30%. However, when considering assistance levels greater than 40%, there is a noticeable difference in the spread or distribution of the projected points compared to the spread observed for assistance levels below 40%.

**Figure 10.**
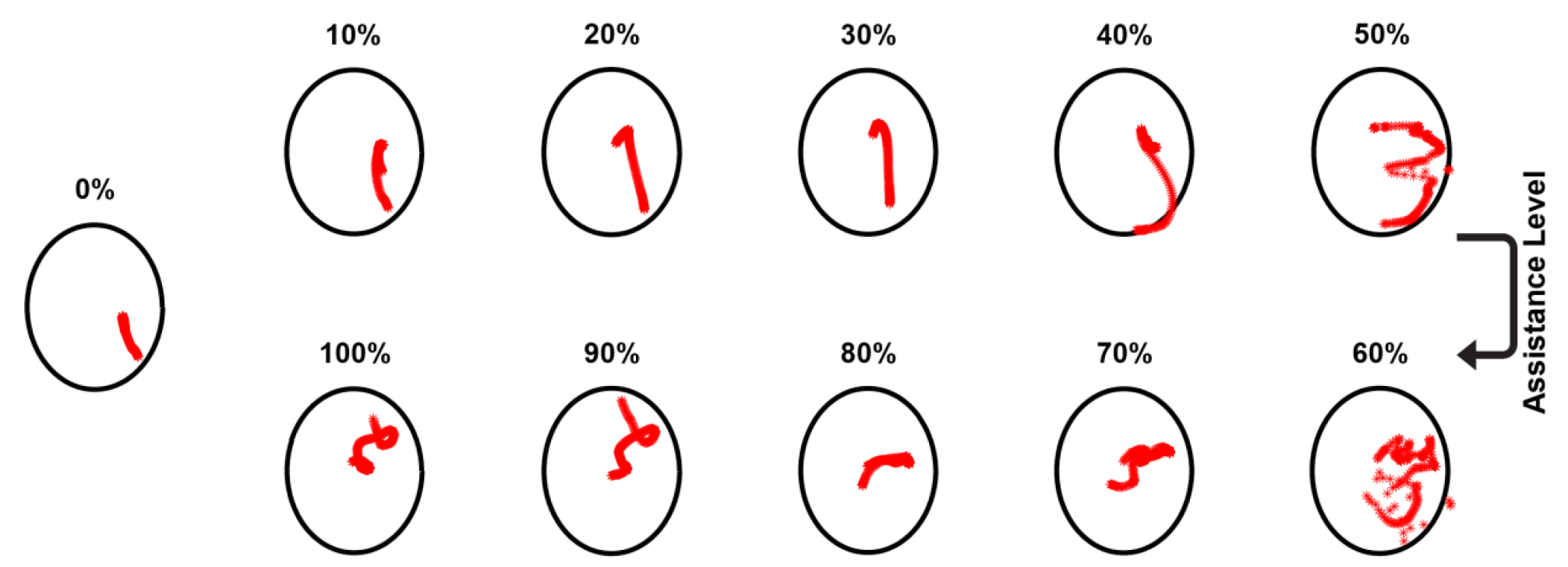
Projection of the glenohumeral contact force of the lateral reaching task onto the glenoid fossa for different assistance levels. The shape of the glenoid fossa was represented as an oval and the projection points were estimated using the ratio of the AP and SI components to the compressive component. The projected points are plotted every 20ms.

## Discussion

Identifying an optimal anti-gravity assistance level or range of levels for passive shoulder exoskeletons can potentially improve muscular efficiency while minimizing the undesirable effects on the user [1, 5, 49]. In this study, we leveraged the task space framework from robotics to perform neuromusculoskeletal modeling and simulations to explore and identify an optimal assistance level for select functional shoulder movements. The simulated dynamic shoulder movements involved elevations in both the coronal and scapular planes, as well as forward and lateral reaching motions. These movements were simulated for two distinct hand target positions, one located at shoulder height and the other above it. Our simulation results demonstrated that, overall, an assistance level above 40% of the maximum gravity torque at the shoulder joint (defined above) can be associated with undesired negative effects during the simulated dynamic arm movements. These included greater end-limb (i.e., hand) movement variability, greater total muscle effort rate, less efficiency, and compromised GH stability and alignment.

The level of anti-gravity assistance applied to the shoulder joint can negatively impact the hand (i.e., end-limb) kinematics and dynamic stability [20]. The results obtained from our predictive task space framework indicate that the ability of the upper limb model to accurately track the desired hand trajectory decreases as the amount of anti-gravity assistive torque at the shoulder joint increases. This eventually led to higher amounts of divergence from the desired trajectory, and therefore hand dynamic instability, when assistance level was above 40%. This observation is consistent with studies on the use of the passive shoulder exoskeletons for overhead tasks, which reported higher errors and reduced quality for tasks that demanded higher precision like overhead drilling [20, 50]. Similarly, studies of robot-assisted rehabilitation for reaching movements reported performance degradation and higher endpoint error with increase in assistive force [51].

The simulated arm movements involved five DOFs of the upper extremity (Table 1), which is greater than the number of DOFs required for the 3-D positioning of the hand in the task space. This kinematic redundancy enables the task-posture controller to effectively decompose the generalized torques into two decoupled torque vectors that correspond to the desired hand trajectory and the arm posture [31]. This decoupling prevents the torques, which are responsible for generating arm postural behaviors, from causing accelerations towards the end-limb task. However, at the muscle level, there are physiological limitations in muscular force generation that may impede the ability of the muscles to meet the kinetic demands of the task-posture control [52]. This potentially explains (1) the arm instability and end-limb divergence from the desired trajectory for high levels of assistive torque and (2) why the onset of hand resultant deviation in 3-D space was primarily coincident with the hand transition from acceleration to deceleration during the advancement phase (Figure 6).

Our results supported our hypothesis that there is an optimal assistance level for which the total shoulder muscle effort rate and efficiency are minimized. Previous studies provide evidence for the existence of an optimal assistance level for a passive ankle exoskeleton based on metabolic cost [53] and for assisted elbow flexion/extension movements based on muscular efficiency [27, 28]. In general, the optimal assistance level appears to be one that balances decreased activations of agonist muscles with increased activations of antagonist muscles. We used total muscle effort rate [22] and efficiency [28] to account for this balance. Our findings suggest that, for most of the simulated dynamic tasks, an assistance level of 10-30% minimized the total muscle effort rate and maximized efficiency. These results are consistent with previous studies that evaluated passive shoulder exoskeletons, such as WPCSE [1] and Exo4Work ]5[ , in which the maximum assistance torque was set to approximately 30% of the maximum shoulder gravity torque.

Interestingly, our results (Figure 8) indicated that, for reaching movements toward a shoulder-height target, an assistance level lower than 30% may be optimal, implying that the optimal level of assistance can be task-dependent. However, the assistance torque provided by some other commercially available passive overhead exoskeletons is closer to 50% [6] of the estimated gravity torque. This discrepancy may be attributed to the fact that commercial exoskeletons are primarily intended for use in static and quasi-static overhead tasks, while our simulated tasks involved dynamic overhead and shoulder-height reaching movements. Another potential explanation for this difference could be related to the fact that many experimental studies primarily employ surface electrodes to measure the electromyographic (EMG) activity of the shoulder muscles. This can limit their assessment of muscular effort to only the superficial muscles and not all the muscles spanning the shoulder joint.

Our study also revealed that high assistance, i.e., beyond 50% for the simulated tasks, might reduce efficiency. This is consistent with previous studies on assistance of single-joint movements, where increasing assistance force beyond an optimal level reduced the efficiency of the muscular response [27, 28, 54]. The provision of shoulder assistance during dynamic or quasi-static tasks can impact the activity of both agonist and antagonist muscle pairs [1, 5, 19, 22]. High levels of gravity-compensating torque have been shown to significantly decrease the activity of shoulder flexor muscles that mainly function to counteract gravity torque (i.e., load bearing muscles). However, the activity of the antagonist muscles, which are essential for joint stabilization and deceleration (i.e., fine-tuning muscles), does not necessarily decrease [36]. Studies have also reported that, in instances where the timing of a perturbing, high force is predictable, individuals tend to voluntarily increase the activation of their muscles in an attempt to stiffen their joint [54, 55]. Therefore, to deliver a higher efficiency, an optimal assistance level should not interfere with the synergistic activation of agonist and antagonist muscle pairs.

Shoulder exoskeletons should be designed to ensure the stability of the GH joint. Our study showed that the glenohumeral joint stability and alignment can be negatively influenced by high levels of shoulder assistance. The GH joint is a ball-and-socket joint that has an inherently unstable structure since the glenoid “socket” does not sufficiently surround the humeral head [56]. The bones, joint capsule , and ligaments contribute most to the static stability of the GH joint [57]. The GH stability during dynamic movements, however, highly depends on the muscles surrounding the joint [57-59]. GH stability is maintained by ensuring that the resultant bone-on-bone joint contact force between the humerus and scapula is oriented within the boundary of the glenoid rim [57]. Our results showed that, with increasing assistance level, the resultant GH joint contact force advanced closer to the edge of the glenoid rim and, therefore, the GH joint became less stable. Such instances of heightened instability and joint misalignment could eventually lead to injury and joint degeneration [60, 61].

Our simulation-based study had some limitations. First, we employed an ideal sinusoidal assistive torque to model the external assistive force/torque of a passive exoskeleton worn by the user. This approach, while simple, allows for generalizability of the results to a range of passive shoulder exoskeletons with differing designs. Nevertheless, in the future, including a model of an exoskeleton with inertial and other dynamic (e.g., damping) properties could provide more insightful and realistic outcomes. A second limitation of our study was that our simulations only explored dynamic reaching and elevating arm movements with fixed initial angles, thereby only capturing a limited subset of possible shoulder movements. To obtain a more comprehensive understanding of user-exoskeleton interaction, future studies should include a larger number of static, quasi-static, and dynamic movements that are representative of the wide range of tasks for which exoskeletons may be used. A third limitation of our study was the use of a fixed movement speed across all simulated tasks. It is important to note that movement speed plays a crucial role in the net torque generated at the shoulder joint during dynamic tasks, thereby affecting the level of muscular activity and demand [50]. Research in gait studies has indicated that the optimal exoskeleton stiffness level (comparable to gravity compensation level) that minimizes steady-state metabolic demand is higher at faster walking speeds [62]. This implies that different optimal assistance levels may be required for movements of varying speeds. Therefore, future studies should consider simulating arm movements at various speeds and explore the potential relationship between the assistance levels and movement speed.

## Conclusion

In summary, the present study demonstrated the feasibility of using a robotics-based task-posture control framework to simulate goal-directed upper limb movements with external assistance. The dynamic task-posture decomposition enabled the execution of the goal-oriented movement in the task space while the assistance was provided in the posture space. Our study suggested that there may be an optimal level of anti-gravity assistance for passive shoulder exoskeletons depending on the types of movements or tasks that they are intended to support. This optimal level of assistance minimizes the user muscular effort rate and achieves a more efficient muscular response without compromising joint stability and limb kinematics. Specifically, our results indicate that for dynamic forward and lateral reaching movements and shoulder elevations in the frontal and scapular planes, an optimal assistance level lies within the range of 20-30% of the maximum gravity torque at the shoulder joint. However, future studies should explore a wider variety of movements with varying speeds to investigate the relationship between the level of assistance and movement speed.

